# Defining the Parameters to Improve Plant Regeneration with Developmental Regulators

**DOI:** 10.1101/2022.11.15.516675

**Authors:** Ryan Nasti, Jon P. Cody, Matt H. Zinselmeier, Nikil B. Badey, Adhvaith Sridhar, Ambika Sharma, Michael F. Maher, Benjamin K. Blackman, Daniel F. Voytas

## Abstract

Tissue culture methods which serve as the standard to regenerate modified plants are challenging and have limited the capacity to engineer new accessions. To improve upon these techniques, genome modifying reagents have been combined with developmental regulators to create gene edited plant tissues in both monocot and eudicot species. Co-culturing seedlings with Agrobacterium strains encoding developmental regulatory genes has proved to be effective at producing *de novo* meristems in multiple eudicot species. In order to see that this technology scales well beyond proof of concept experiments, various parameters were tested for refinement. Improvements have been observed at the key stages of growth induction and progression to shooting by manipulating the vector design, developmental regulator choice, Agrobacterium strain selection, and regulatory gene removal systems. Having defined these parameters as viable optimization points, the avenues to apply developmental regulators for plant regeneration in more diverse species have become more feasible.

## Introduction

Conventional methods to genetically engineer plants most commonly utilize *Agrobacterium tumefaciens* or biolistic bombardment to deliver transgenes into host cells. The transfected cells are selected and regenerated into whole plants through rounds of sequential hormone treatments. Despite advances aimed at improving the efficacy of these techniques, regenerating from transformed explants is a major hurdle limiting the creation of new plant accessions.^1,2^ One avenue to overcoming these bottlenecks has been demonstrated with the ectopic overexpression of developmental regulatory genes (DRs) that control specific organ inducing pathways.^3–5^ Combinations of DRs such as *Zea mays Wuschel2* (*ZmWus2*) and *Baby Boom* (*Bbm*), promote production of transgenic or gene edited somatic embryos across diverse monocot species.^6,7^ In eudicot species, the application of *ZmWUS2* and either *SHOOTMERISTEMLESS* (*STM*) or *isopentyl transferase* (*ipt*) leads to the formation of *de novo* meristems.^5^ Additional wound-responsive DRs, in particular *PLETHORA* (*PLT5*), have also been shown to promote meristem formation in multiple eudicot species.^8^ The accumulation of these different DR-based approaches promises increased ease and scalability for the generation of both transgenic and gene edited plant lines.^9^

While co-opting organogenesis for gene editing has proved effective, there still remains further room for optimization towards the goal of increased scalability. If we look specifically at the development of *de novo* meristems in eudicots, transfected tissues transition first to globular growths, then to shoots and eventually into plantlets.^10^ Within this progression, the total growth formation and the downstream shooting frequency serve as the primary transition steps (Figure 1). Testing parameters to increase the efficiency of each of these steps in *Nicotiana benthamiana* would yield a higher overall capacity to generate modified plantlets. Insights found from these tests in this model system can then be used to inform the development of similar methods in other plant systems.

**Figure 1.**
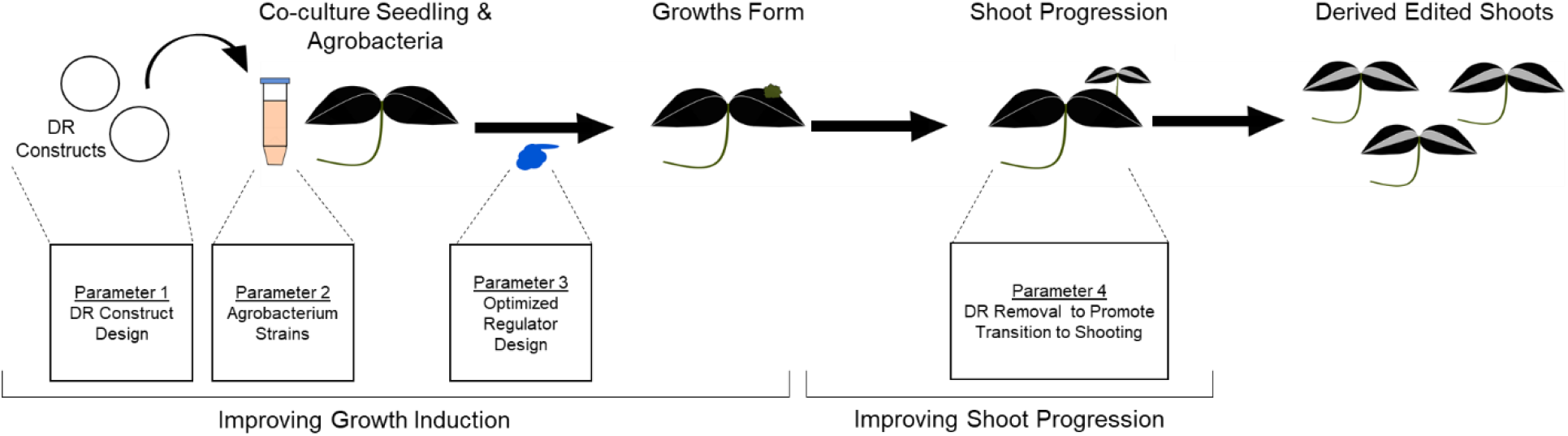
Optimization Parameters for DR-Based Transformation. During the process of *de novo* meristem induction after Fast-TrACC co-culture, there are two pivotal transition points: growth induction and shoot progression. For each of these transition points there are key parameters to optimize, namely construct design, use of different Agrobacterium strains, use of alternate DR variants, and controlling DR removal. By making alterations to these parameters the overall efficiency of DR-based regeneration can be improved.

Total growth production proves to be a particularly vital step since the number of regenerable plants derives entirely from the initial pool of formed growths. Therefore in order to see improvement in this area, it is pertinent to assess a diverse set of parameters for their impact on growth induction. A foundational parameter for growth induction is the specific design of the delivery vectors for the DRs. In monocots, there is evidence that increases in embryogenesis can occur from either high levels of localized expression or by splitting delivery of DRs onto two separate Agrobacterium strains.^11^ Testing principles like these in *N. benthamiana* can lead to similar improvements for *de novo* meristems.

Another area for growth improvement manifests with the DRs themselves. While developmental transcription factors tend to be highly conserved across groups of plant species,^12^ DRs from more closely related species could hold the potential to increase growth yields. Taking this principle further, synthetically engineering one or more of these DRs could improve transformation efficiency. Of the DRs that have been tested, the *WUS* transcription factor is a common feature across DR technologies and thus serves as an ideal candidate for engineering. In its native context, *WUS* serves the pivotal role of promoting cell division within the organizing center of the meristem.^13^ Rationally altering less essential functional domains provides the potential to improve the efficacy of *WUS* for biotechnological applications.

A final factor for the improvement of growth formation comes by comparing Agrobacterium strains with differing delivery potentials. Agrobacterium serves as the fundamental component in the Fast Treated Agrobacterium Co-Culture (Fast-TrACC) method, as a highly virulent Agrobacterium strain allows for more efficient T-DNA delivery.^14^ Interestingly, the T-DNA transfer capability of different strains can vary widely even if derived from the same parent strain (Supplemental Table 2).^15^ Finding the Agrobacterium strain with the optimal delivery pattern for the target tissue of a given species would improve downstream growth production from the DRs.

In addition to growth induction, alterations to shoot progression will also have an important impact on the efficiency of plant generation. For proper shoot progression, one factor preventing this transition is the continued presence of the DRs. Since one of the primary functions of DR expression is to promote cell division, the transition into normal shoot development could be hindered by their maintained expression.^3,5,6^ Certain growth promoting DRs, such as *GRF/GIF* or *PLT5*, have been shown to promote the formation of shoots from callus, but have yet to be demonstrated in combination with additional DRs.^8,16,17^ Their incorporation with division inducing DRs, like *ZmWUS2*, could be one avenue to trigger the transition to shooting. In monocot systems, defining additional ways to overcome the cell division programming has been important for the application of DRs. By using techniques involving DR excision or spatial DR separation the undifferentiated state was able to be overcome and shoots could form.^3,11^ Taking such practices and applying them for DR use in *N. benthamiana*, we will gain new insights into how the transition to shooting can be improved.

To these ends, we herein describe efforts to test these different parameters and clearly establish the principles required to improve the efficiency of DR methods. These comparisons allowed us to define the essential tenets that were critical in *N. benthamiana* that should be tested for use in across species in subsequent DR-based plant transformation approaches.

## Methods

### Construct Design

Constructs described herein were assembled using the previously published golden gate based modular cloning platform.^18^ With layered type IIS restriction enzymes, sequences (individual sequence design and acquisition detailed in the relevant sections) were first cloned into individual modules, and subsequently into the applicable T-DNA vectors. A multitude of constructs were created to contain different combinations of DRs and reporters (Supplemental Table 1). These plasmids and their related DNA sequences will be made available at Addgene. These T-DNA vectors were subsequently transformed into Agrobacterium strains for use in co-culture. The specific Agrobacterium strains used are detailed for each of the individual experiments.

### General Fast-TrACC Reagent Delivery & Growth Assessment Procedure

DR constructs were delivered to *N. Benthamiana* seedlings using Fast-TrACC.^5,14^ This involves transforming the DR construct into your Agrobacterium strain of choice (unless otherwise stated the predominantly used strains were: GV3101 for *WUS* variant experiments and C58C1 for both CRE-lox and vector design experiments) and co-culturing this Agrobacterium with *N. benthamiana* seedlings germinated in liquid culture.^10^ Two days after being removed from co-culture, seedlings are assessed for reporter signal to determine successful T-DNA transfer. For the constructs encoding luciferase, plates containing greater than 67% luminescent seedlings showing luciferase positivity were tracked for growth formation.

Growths typically begin forming within the window of 10-14 post co-cultivation, but can vary depending on chamber light conditions. At this stage any observable growths will be visible but are likely small (Supplemental Figure 1, purple arrows) and harder to count. At 20 days post co-cultivation, the growths should be clearly discernible for counting (Supplemental Figure 1). Individual growths (Supplemental Figure 1, arrows) are simpler to count than clusters of many growths (Supplemental Figure 1, white circles). For clustered growths best estimates were made based on the appearance of darker green puncta within the grouping. Despite efforts to keep growth formation as consistent as possible, delivery and subsequent transformation are highly variable across experimental replicates. For this reason, total numbers of induced growths can range quite strongly depending on a given experiment. Luckily, the variability is consistent across treatments within a given experiment (the lowest replicate for one treatment tracks with the lowest in the others) and thus the general trends can still be understood.

The components for the majority of the experiments were encoded on a single T-DNA vector. The experiments with pooled vectors are performed with transformed Agrobacterium strains each containing a different T-DNA vector. The strains are then mixed with defined percentages of each strain (Figure 1b-c, for 1b the split ratio is 1:1). To ensure as consistent of DR ratios as possible, the OD600 of the individual strains is normalized to approximately 0.14 before being combined.

### Sequence Comparison and Synteny Analyses

For the cross species *WUS* comparisons, the genomic sequence was isolated for each species’ (tomato, potato and grape) closest homolog to *AtWUS*. This was determined by performing a BLAST search against each representative genome and selecting the gene with the most matched sequence (*SlWUS*: Solyc02g083950, *StWUS*: PGSC0003DMT400009157, *VvWUS*: VIT_04s0023g03310). These sequences were then PCR amplified and incorporated into plasmid backbones for downstream golden gate cloning.

In order to compare a wider diversity of *WUS* sequence variants from across species individual coding sequences were isolated from three *Brassicaceous* species (*Brassica napus*, *Brassica oleracea*, *Brassica rapa*) as well as cassava, cotton and sunflower (*Manihot esculenta*, *Gossypium hirsutum*, *Helianthus annuus* respectively). Similarly to the aforementioned cloned *WUS* sequences, the additional species *WUS* coding sequences were isolated after performing BLAST on the relevant genomes. Several of these species possess multiple *WUS* homologs all of which were used for the analysis. The comparisons made within this study are less comprehensive than other works^12^ and are more targeted for our biotechnological applications. Synteny was assessed using the BioConductR package ‘DECIPHER’.

When cloning the *AtWUS* deletion variants, only the coding sequence was selected for alteration. Using the defined sequence motifs as described in “Rodriguez et al 2016 PNAS” the *WOX* homeobox, the homodimerization domain, the *HAM* binding domain, the *WUS* box & the *EAR*-like domains were all able to be annotated. Based on their previously described functions three deletions were created removing all domains other than the *WOX* homeobox (*AtWUS-D1*), removing the homodimerization domain (*AtWUS-D2*), and removing the *WOX* homeobox and homodimerization domains (*AtWUS-D3*) (Figure 3c). These were meant to emulate non-activating mutants, non-complexing mutants and non-binding mutants, respectively.^19^

**Figure 2.**
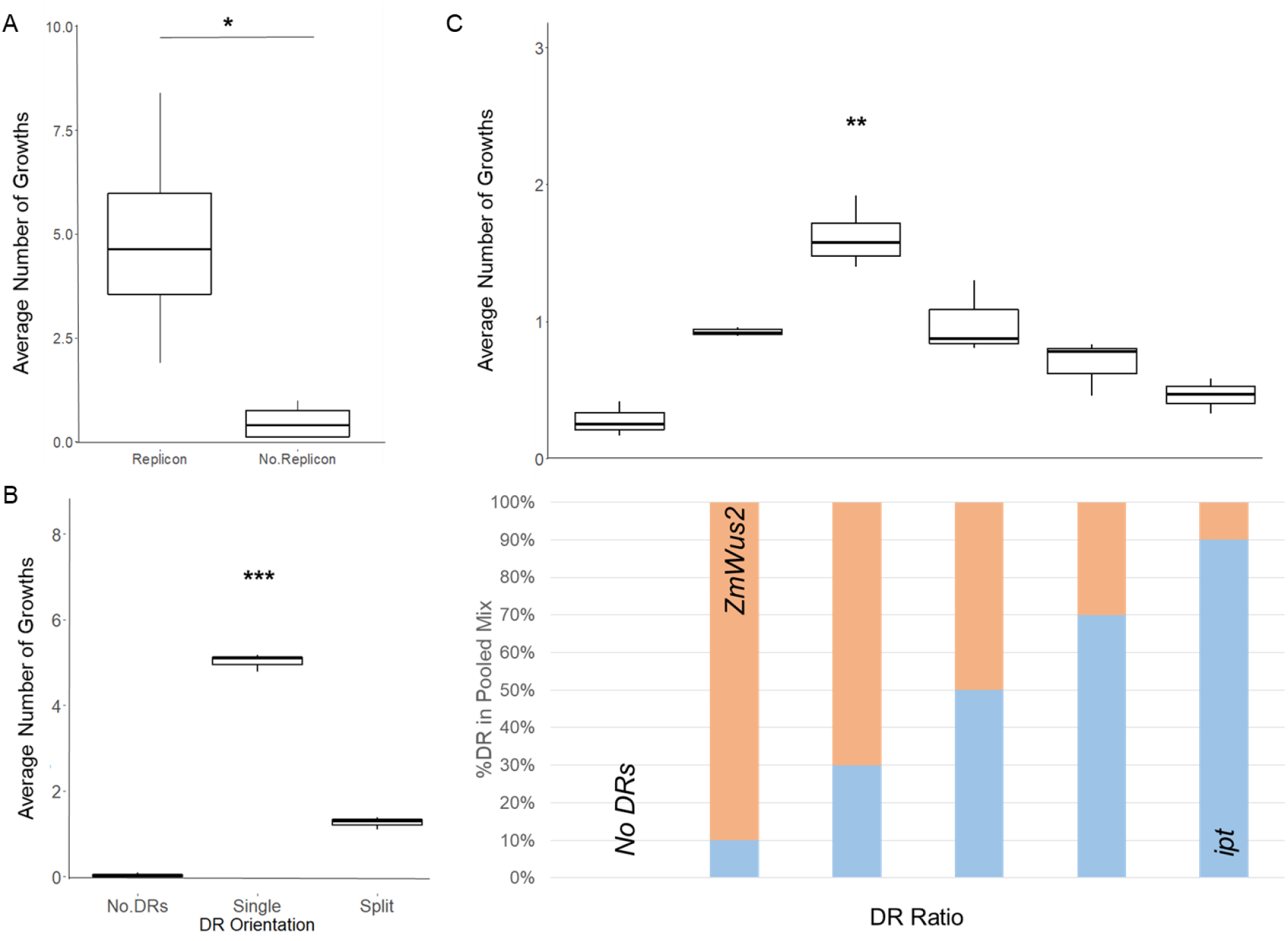
Vector Optimizations for Increased Growth Induction. In order to improve the growth induction efficiency of Fast-TrACC delivery of DRs, several vector parameters were tested to determine their impact on the process. Delivery of Agrobacterium strains with either a T-DNA with or without the components to produce a GVR showed that the GVR’s inclusion increases growth induction (a). Delivery of DR vectors in *trans* has previously been shown to increase growth formation potential in other species, but when applied in *N. benthamiana* the *trans* orientation produces significantly lower growth potential (b). To determine if ratio of the two DRs in *trans* orientation has an impact on growth induction, the ratio of *ZmWUS2* and *ipt* constructs were varied (c), which resulted in the most growths with 70% *ZmWUS2* and 30% *ipt*.

**Figure 3.**
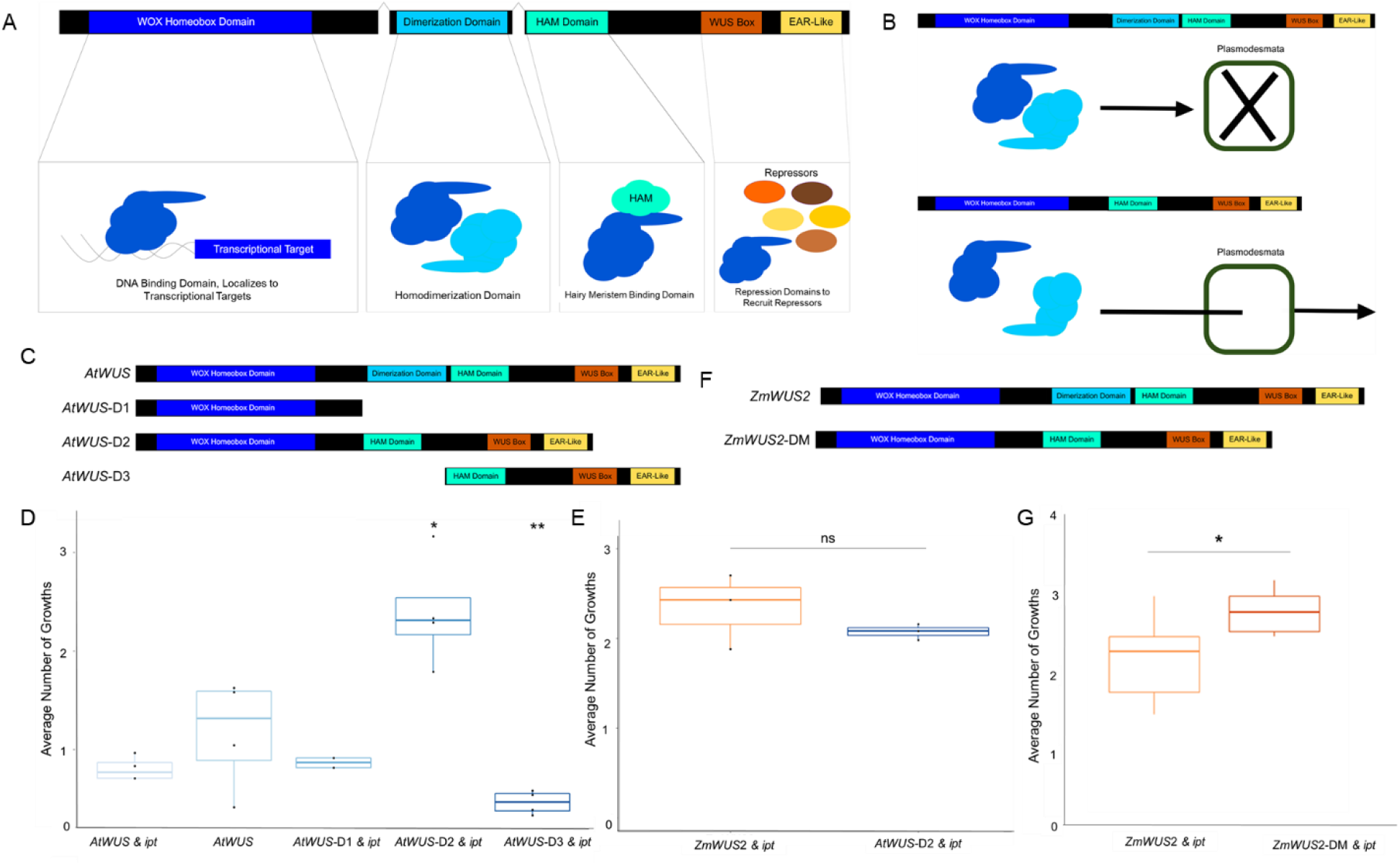
Synthetically Altering the *WUS* Homodimerization Domain to Improve Growth Formation. The transcription factor *WUS* has key functional domains that define how the protein is regulated (a). The WOX Homeobox domain is required for binding at the genomic target sites for regulation. The homodimerization domain and HAIRY MERISTEM (HAM) are involved in binding regulatory co-factors. Lastly the WUS box and EAR-like domains recruit co-repressors to enable *WUS’* transcriptional control of its genomic targets that lead to cell division within the meristem. The homodimerization domain of *WUS* plays the functional role of sequestering its mobility. With a functional dimerization domain *WUS* will bind with another *WUS* blocking its passive diffusion through the plasmodesmata (b, top), but when removed this binding is blocked and the smaller *WUS* is able to move between cells (b, bottom). In order to compare the effects of the different domains of *AtWUS* on its growth induction potential, three deletion mutants were created (c) and delivered in tandem with *ipt*. No significant difference in growth induction was seen between *AtWUS* with or without *ipt* and *AtWUS*-D1 and *ipt* (d). A significant decrease in growth induction to near zero was seen when DNA binding is removed as in *AtWUS*-D3. The only deletion variant that displayed a significant growth increase was *AtWUS*-D2, which removed the homodimerization domain. The *AtWUS*-D2 variant even showed improvement to the point of having no significant difference when compared to *ZmWUS2* (e). Using this as the basis for *ZmWUS2* alteration, the putative homodimerization was deleted from its sequence creating *ZmWUS2*-DM (f). When delivered with *ipt* the number of growths induced by *ZmWUS2*-DM is significantly increased over those from *ZmWUS2* and *ipt* (g).

Looking to extend the domain deletion principle to *ZmWUS2*, its dimerization domain first needed to be annotated. While the *WUS* dimerization domain for *A. thaliana* has been annotated,^19^ this domain has yet to be defined for monocot species. To determine the most likely sequence comprising the dimerization domain for *ZmWUS2*, we produced a series of sequence alignments (Supplemental Figure 4). First, the *WUS* sequences from dicot species were aligned, namely *AtWUS*, *SlWUS*, and *VvWUS*. We utilized the previously annotated *A. thaliana* dimerization domain to define the boundaries for a putative consensus dicot *WUS* dimerization domain. In this alignment, the sequences comprising the *WOX* homeobox, the *WUS* box & the *EAR*-like motif showed high levels of conservation as previously demonstrated.^12^ This analysis identified additional conserved residues located between the *WOX* homeobox and *WUS* box, including a conserved glycine-aromatic patch (Supplemental Figure 4). This sequence could play a role in *WUS* dimerization due to the importance of glycine-aromatic patches in protein-protein interactions and aggregation processes.^20^ Thus, we reasoned the intervening sequence between these N and C terminal domains likely represented the dimerization and *HAM* domains across species (Figure 3, Supplemental Figure 4).

Using this putative dicot dimerization domain for reference another *WUS* sequence alignment was performed with *Wus2* variants from two monocots, *Zea mays* and *Sorghum bicolor*. We again noticed a glycine-aromatic patch within the monocot *WUS* sequences at a similar location within the consensus dimerization domain (Supplemental Figure 4). With these alignments, we reasoned that the glycine-aromatic patch may be an indicator of the dimerization domain’s location for a given *WUS* protein.

### RUBY Image Analysis

Pictures of whole plates were taken seven days after removing the seedlings from co-culture (Figure 4a, Supplemental Figure 6a). At this point a reliable level of RUBY pigment will have accumulated, which should roughly match the peak expression level. The plates were cultivated for another 14 days after which growth progression was assessed. This is consistent with growth counts for the other experiments.

**Figure 4.**
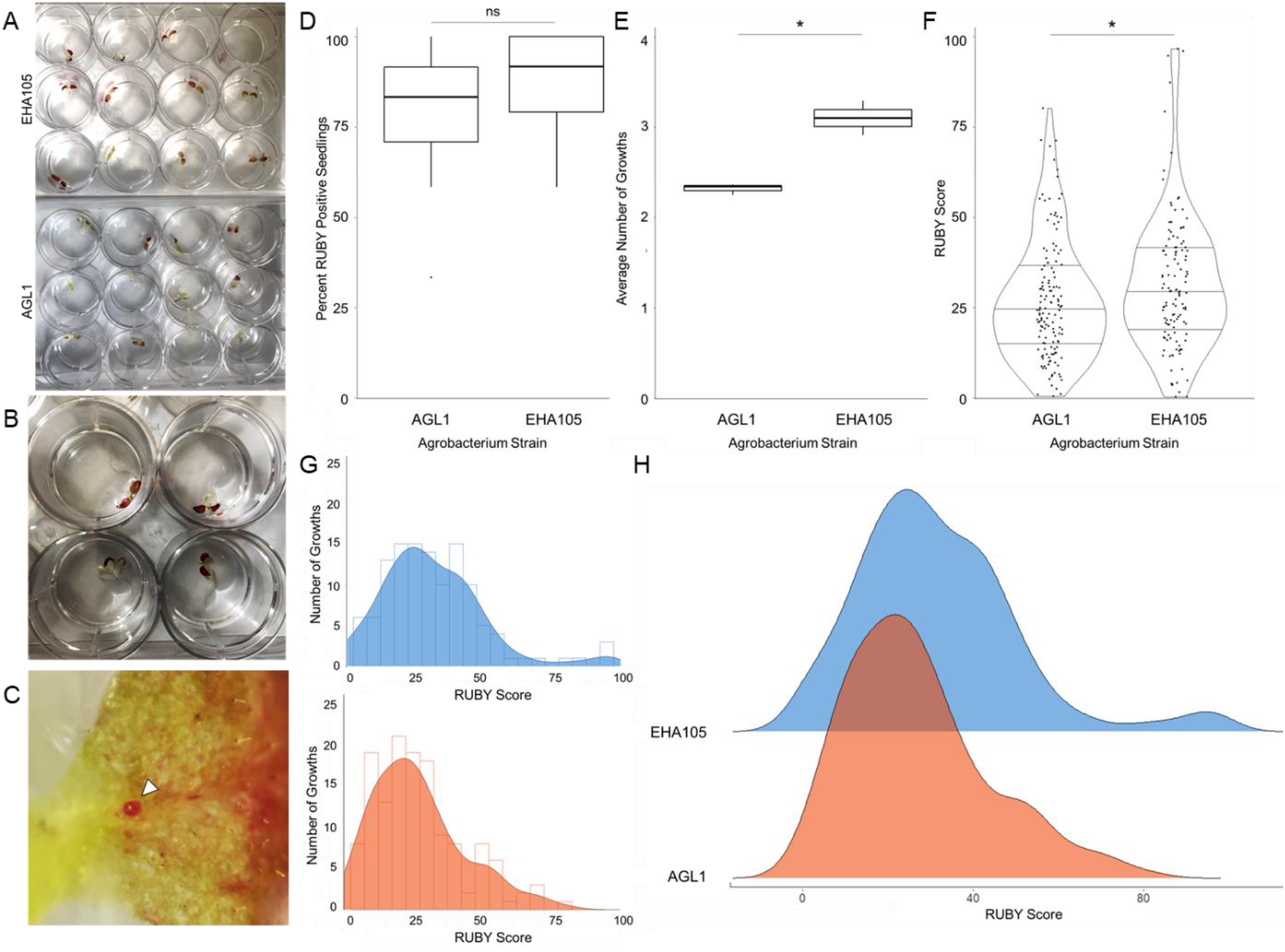
Efficiency of Reagent Delivery from Agrobacterium Determines the Potential for Growth Induction. To evaluate the impact of different two Agrobacterium strains, EHA105 and AGL1 (a), on growth induction two metrics were used: delivery of the RUBY reporter (b) and counts of positive growths (c). After co-culture with the Agrobacterium strains, it was observed that the total number of seedlings with reporter signal was not significantly different between the two (d). Despite delivery to a similar number of seedlings, the number of growths induced from the seedlings treated with EHA105 was significantly higher than those treated with AGL1 (e). This in part is due to the difference in the total area of each seedling that was transfected, with the EHA105 treated seedlings manifesting with a significantly greater RUBY posited sectors than AGL1 (f). The delineation in growth potential manifests in the middle delivery range (g), wherein EHA105 (orange) has a higher number of those middle transformed seedlings than AGL1 (blue) resulting in a higher growth count. There is a diminishing effect of higher reporter positive area as growth induction with either strain begins to decrease after an optimal delivery value (h).

Analysis of red pigmentation intensity in the images of RUBY treated seedlings was optimized by converting two dimensional plant-tissue images into three dimensional intensity plots (Supplemental Figure 6c). A .jpg or .png was first uploaded to ImageJ Fiji software and applied to a pipeline to create the surface plot. Selections from this image may be chosen for exclusive analysis. Since ideal plotting results are achieved when the image dimensions are equal to those of the plot grid, the size of the plot grid is redefined to align with the size of the uploaded picture. Image noise is then smoothed to allow unobstructed measurement of the original object’s characteristics. The lighting of the image is adjusted to enhance the visibility of small differences between adjacent pixels. After the pre-analysis image improvements are executed, the image is then analyzed as a combination of pixels wherein the plane of the image is used to internally define an x and y direction. The color is measured per pixel, with each pixel being assigned a numerical value for the representative red coloration intensity. This value is then appended to the coordinate location (x, y) as an additional z-coordinate to be plotted as height. Longer visual spikes represent an increased concentration of red for the respective pixel.

In addition to the whole plate signal intensity analysis, the RUBY coverage for each seedling was also assessed. From the whole plate images, each of the wells within were isolated for individual seedling analysis. From these individual image partitions, the collection of RUBY positive pixels was divided by the pixels encompassing the total area of the seedling to give a RUBY score. These RUBY scores were then correlated with the total number of growths on that same individual seedling (Supplemental Figure 7). To define a general trend for the data growth versus RUBY score, the data was run through a loess-based smoothing statistical analysis (Figure 4f-g).

### Monitoring Shoot Progression after CLV3::CRE Treatment

Similarly as with tracking growth induction and progression, *N. benthamiana* seedlings were grown for 15-20 days after co-culture before making observations. From cotyledon tissues with any noticeable mass of growths, those sections were excised (not separated) and transferred to ½ MS media plates. Once on these plated the explants were monitored for the progression of any differentiated tissues (e.g. leaflets, shoots, etc.). At 30 days post transfer to plates (50-60 after co-culture), shoot numbers were collected to be compared between the *CLV3*::*CRE* and No *CRE* treated tissues. Explants derived from the treatment without *CRE* did not look appreciably different from those treated with *CLV3*::*CRE* before they transition to shoots despite their higher transition rate.

## Results

The implementation of DRs to facilitate regeneration has been demonstrated in several species.^3–5,16,21–23^ Specifically for *de novo* meristem induction, the majority of examples of DR application were performed in *N. benthamiana*.^5^ Despite the observed utility of both direct injection and Fast-TrACC methods to produce meristems, a multitude of components can be further improved. Focusing on the Fast-TrACC method, four parameters were targeted for improvement (Figure 1): vector design, DR choices, the use of highly virulent Agrobacterium strains, and controlled removal of DR expression.

### DR Vector Design Parameters

Previous Fast-TrACC regeneration experiments employed T-DNAs encoding both DRs as well as the components for a geminiviral replicon (GVR).^5^ These replicons have been shown to substantially increase the expression level from a given T-DNA.^24^ Therefore components encoded within the replicon would theoretically produce higher levels of the DRs than without a replicon. To test experimentally whether or not these GVR components were truly necessary, we compared the rate of growth induction from T-DNA cassettes encoding the DRs with and without the replicon motifs. Not only was the replicon cassette more efficient at inducing growth formation, but the T-DNA without the replicon components promoted effectively no growths (Figure 2a). This indicated to us that the GVR was necessary for growth formation in this system.

In addition to the expression level increase provided by the GVRs, there is increasing evidence that delivering the DRs on separate vectors can have a positive effect on regeneration potential.^11^ This trend is likely due to the cell to cell mobility of *WUS*.^19^ While such results are promising in these other species, this delivery schema has yet to be tested in *N. benthamiana*. To this end, Agrobacterium strains containing *ZmWUS2* or *ipt* respectively were pooled at a 1:1 ratio and co-cultured with *N. benthamiana* seedlings. After Fast-TrACC, growths were counted and contrary to other systems the rate of induction is lower in the *trans* orientation than the *cis* (Figure 2b). This suggests a potential species or tissue specificity exists as *trans* DR delivery has yielded increased growth rate in other species.^11^

In monocot systems where *trans* delivery of DRs was successful, different stoichiometric ratios of delivered regulators yielded differing rates of organogenesis. Even with a decreased amount of *ZmWUS2* compared to *BBM*, a transgenic somatic embryos were still able to be readily created.^11^ Building upon this principle and applying it to *de novo* meristem induction in eudicots, we pooled *ZmWUS2* and *ipt* at ratios of 9:1, 7:3 and 1:1 to determine if the ratio impacts growth formation (Figure 2c, bottom). From these different DR ratios, a negative parabolic relationship was observed, with the 70% *ZmWus2* pooled treatment showing the highest growth induction potential (Figure 2c, top). While the number of growths is lower in this schema, it is useful to understand how it functions as it could be used in tandem with an additional DR-free trait vector. As previously mentioned, *cis* and *trans* DR delivery orientations have differential capacity across species and potentially even tissue types. Therefore it is important to define the best set of parameters for your individual system when implementing such techniques.

### Testing Different *WUS* Variants for Growth Induction Potential

Of the different DRs that have been implemented for *de novo* organogenesis, the *WUS* transcription factor has been pivotal across plant transformation applications. To this point, the *Zea mays Wus2* (*ZmWus2*) variant of *WUS* has been predominantly used in these applications. This variant has been shown to be effective in species ranging from monocots like maize and sorghum, to eudicots including *N. benthamiana*, and tomato.^4–6^ These species cover a wide evolutionary range, yet the same *WUS* variant is able to function effectively. This phenomenon is particularly striking for the dicot species where the level of synteny between the *ZmWUS2* variant and endogenous *WUS* is mostly limited to the *WOX* homeobox domain (Supplemental Figure 2d & 4). This begs the question as to whether the *WUS’* origin matters when applying to increasingly diverse groups of species.

To determine if the species of origin has an impact on a *WUS* variant’s ability to induce *de novo* growth formation, *WUS* sequences from four eudicot species were selected to be compared to the *ZmWUS2*, namely tomato, potato and grape and *Arabidopsis thaliana* (*SlWUS*, *StWUS*, *VvWUS*, *AtWUS* respectively). There was no significant difference observed between the growth induction potentials of *VvWUS* or *SlWUS* and *ZmWUS2* (Supplemental Figure 2a). Growth counts after Fast-TrACC delivery of the *AtWUS* coding sequence were also significantly lower than growth counts from *ZmWUS2* (Figure 3b). *AtWUS* similar to the coding sequences of other *Brassicaceous WUS* varieties, phylogenetically clusters separately from other examined eudicot *WUS* variants (Supplemental Figure 3a), likely due to a conserved sequence motif found specifically in these species (Supplemental Figure 3b). This unique sequence motif likely has an impact on the effectiveness of *AtWUS* to induce growths in *N. benthamiana*. Despite these few outlier sequences, the majority of tested dicot *WUS* variants function similarly to *ZmWUS2* indicating that species of origin plays a minor role in the ability of a *WUS* variant to produce *de novo* growths.

Despite the lowered function of *AtWUS*, it contains well-studied functional domains (Figure 3a). Through knock-out studies in *A. thaliana*, certain domains have been described either as essential more plastic in *WUS’* function.^19^ One such domain is the homodimerization domain, which has been shown to regulate *WUS* movement between cells. When two *WUS* proteins bind one another they form a larger protein complex that can not passively diffuse through the plasmodesmata. Removing this domain should thus allow for the smaller *WUS* proteins to move into neighboring cells and impact a greater proportion of the treated tissue (Figure 3b). To test this hypothesis, three deletion mutants (Figure 3c) were made to the coding sequence of *AtWUS*. Testing against the growth forming capacity of *AtWUS* & *ipt* or *AtWUS* alone, no significant change was seen delivering *AtWUS-D1* and a significant decrease was seen with *AtWUS-D3* (Figure 3d). Of the three the only variant to show improvement was *AtWUS-D2* which lacked the homodimerization domain (Figure 3d). *AtWUS-D2* was even able to rescue its function in *N. benthamiana*, showing no significant difference when compared to *ZmWUS2* (Figure 3e). With the enhanced function of *AtWUS-D2*, the capacity for engineering of *WUS* variants has been established.

Taking this principle of homodimerization domain removal further and applying it to the more functional maize variant should improve its ability to induce growths. Improvement to this variant will be particularly useful due to its usage in both monocots and eudicots.^5,6^ Thus, we generated a *ZmWUS2* truncation to remove the glycine-aromatic patch to create a *ZmWUS2* dimerization domain mutant (*ZmWUS2*-DM) (Figure 3f, Supplemental Figure 4). This deletion variant is again expected to improve *ZmWUS2*’s capacity for passive diffusion^19^ and subsequently leading to increased growth formation. The *ZmWUS2*-DM and *ipt* combination significantly improved increased growth counts when compared to full length *ZmWUS2* and *ipt* (Figure 3g), albeit very modestly. In addition, the *ZmWUS2-DM* treatment has the capacity to intermittently create stretches of amorphous growth along the cotyledon edge not typically seen with *ZmWUS2* (Supplemental Figure 5). This patterning makes sense with the presumed loss of spatial patterning conferred with the homodimerization domain. In the case of *ZmWUS2*, the growth induction potential is likely near its maximum, thus modifications yield only minor improvements to its effectivity. As it stands *ZmWUS2* is a sufficient regeneration tool without the alterations made to *ZmWUS2-DM*. However, the engineering principle displayed with both *AtWUS*-D2 and *ZmWUS2-DM* illustrate avenues to enhance species specific *WUS* variants. Therefore these improvements in growth induction demonstrate the impacts of rational DR engineering to increase transformation efficiency.

### Impact of Agrobacterium Strain on Successful Growth Induction

Since Agrobacterium is essential to Fast-TrACC delivery of the DRs with Fast-TrACC, understanding the transfer capacity of different strains and the downstream growth potential will prove important for further optimization. Previous research has established the importance of highly virulent Agrobacterium strains for *de novo* organogenesis to occur effectively.^3^ In order to determine the impact of Agrobacterium on the Fast-TrACC method, a collection of strains from different sources were compared against one another. Across these strains, a high amount of variability in growth induction was observed (Supplemental Table 2). This variation was even seen in different variants of a given parental strain, such as GV3101.

Looking to further define what underlies this variation, two strains were selected for further examination, namely EHA105 and AGL1. To quantify each strain’s extent of delivery, the RUBY reporter was employed to track with a clear visual phenotype.^25^ The two strains were transformed with a construct encoding *CmYLCV*::RUBY along with the DRs *ZmWUS2* & *ipt* and subsequently co-cultured with *N. benthamiana* seedlings. Where Fast-TrACC delivery was successful deep red pigmentation is observed (Figure 4a-b, Supplemental Figure 6a-b). By imaging RUBY positive seedlings and tracking for growths (Figure 4c) we can link total delivery with induction potential. Interestingly, neither Agrobacterium strain showed a significant difference in the total number of transfected seedlings (Figure 4d) despite showing altered growth induction (Figure 4e). The key difference appeared in the distribution and intensity of the RUBY signal, with EHA105 transfecting a larger more strongly pigmented areas than AGL1 (Figure 4f, Supplemental Figure 6c). This differential reagent transfer underlies the significant increase of growths. This becomes particularly apparent in the middle range of RUBY scores (~30-60) where the EHA105 growth trend (Figure 4g, i: blue) shows a higher abundance of individuals producing growths compared to AGL1 (Figure 4h, i: orange). Interestingly, for either Agrobacterium strain there is a maxima at which growths optimally seem to appear, wherein too much DR expression begins to have a detrimental impact on growth formation (Figure 4g-h). Since the RUBY reporter is known to shunt out of tyrosine biosynthesis, it needed to be determined if this drop off was due to RUBY or DR expression. By performing leaf infiltrations with different DR combinations, DRs were found to cause tissue death when expressed at higher levels (Supplemental Figure 8). These reagent delivery trends highlight the importance of selecting an Agrobacterium strain with optimal properties for a given species of interest.

### DR Excision with Recombinases

Even with the observed capacity of DRs to induce growths, progression to shoots from these growths is dependent on overcoming the patterning caused by these DRs. Since *WUS* strongly promotes cell division within the meristem,^13^ the majority of growths that form do not properly progress to shoots.^5^ Additionally, maintained DR expression has the potential to impart strong developmental dysregulation on plants regenerated in this fashion.^3,5^ Due to these potential long-term developmental defects, applications in monocots were implemented in tandem with techniques to control DR expression. Whether tight tissue specific expression or DR removal these methods have been shown to remove dysregulation and improve regeneration efficiencies.^3,6^ One particular tool used for DR removal is the CRE recombinase. Recombinases like CRE recognize specific DNA motifs and facilitate the exchange of sequences between them.^26^ By combining the principles of tissue specific expression and DR excision there exists the potential to discretely control when and where the DRs are expressed. With this logic in mind, we sought to synthetically build an alternative, self-contained *WUS* regulatory network.

Endogenously in the meristem, *WUS* will transcriptionally activate *CLV3* which begins a cascade to downregulate its own expression (Figure 5a).^13^ This feedback loop provides the balance between cell division and differentiation necessary for functional *in planta* meristem progression. With our ectopic overexpression, the transfected *ZmWUS2* will not be deactivated in the same capacity as the endogenous *WUS*. Due to this, some portion of the transfected cells will continue their cell division unchecked and will not begin to differentiate. Looking to combat this continuous developmental impact, a sought to devise a CRE recombinase removal system to excise the *ZmWUS2* coding sequence whenever it is highly expressed (Figure 5a).. Due to *WUS’s* capacity to bind the *CLV3* promoter, when this promoter is used to express CRE it should be tightly linked to where *ZmWUS2* is being overexpressed. With the initial cell division programming sufficiently in place, removing the DRs at this point should allow for the transition into more shoot-like tissues. When comparing the *CLV3*::CRE removal system with the standard DR delivery platform, even though the total number of growth sections that are produced is higher with no CRE expression (Figure 5b-c) a substantially greater amount of shooting is seen when CRE excises the DRs (Figure 5b-d). This builds credence to the idea that excess *WUS* expression is limiting the amount of shooting occurring, with a multitude of shoots occurring at sites of assumed excision (Figure 5e).

**Figure 5.**
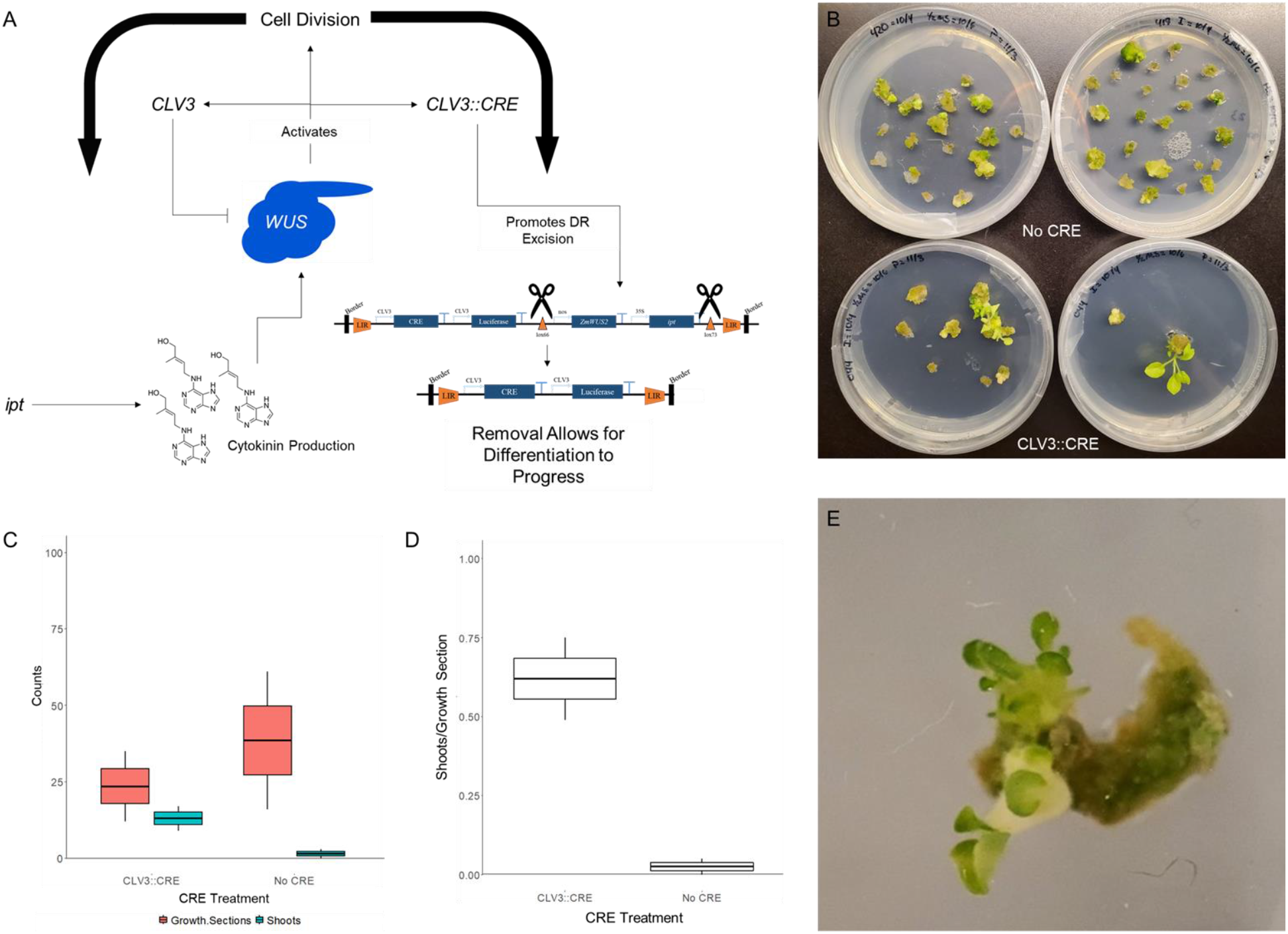
Employing CRE Recombinases to Promote DR Excision and Subsequent Shooting. In the endogenous meristem, *WUS* and *CLV3* establish a feedback loop to control the expression levels of *WUS*. Taking advantage of this regulatory loop, the *CLV3* promoter was used to express CRE and subsequently build a synthetic DR removal platform (a). By using *WUS’* inherent ability to bind and activate at the *CLV3* promoter, when *WUS* is expressed in the presence of *CLV3*::CRE it will excise its own expression cassette. This synthetic circuit therefore recapitulates the endogenous control mechanism. Tracking growth sections from seedlings treated either without CRE or with the *CLV3*::CRE. The CRE free treatment results in more growths overall (b top & c), but in the growths derived from the CLV3::CRE treatment shooting was more abundant (b bottom & c-d). When this successful shooting occurs, there are often multiple shooting events from a given site (e).

## Discussion

To overcome the difficulties present in the field of plant transformation, DRs have been used to improve the ease and universality of regenerating both eudicot and monocot species.^3–5,16,21–23^ Despite the promise of their application, these technologies are early in their development. Improving different components within these methods has the potential to further impact the efficiency of DR-based transformation towards plant transformation at scale.^9^ Using the Fast-TrACC delivery method in *N. benthamiana* we have a rapid and comparable platform to define and optimize the factors most strongly impacting the capacity to generate new plant accessions with DRs.

By focusing on the steps of growth induction and shoot progression we can target two of the major transition steps in the production of these *de novo* growths. In *N. benthamiana*, we have established that the highest levels of growths can be induced when the DRs *ZmWUS2* and *ipt* are delivered on the same vector with a highly virulent Agrobacterium strain (Fig 2 & 4). Taking this principle a step further and synthetically modifying *WUS* can also improve growth induction while potentially removing the counteracting DR persistence (Figure 3). Prolonged DR expression serves as the main limitation in shoot progression. By designing synthetic removal systems, whether chemically or transcriptionally induced (Figure 5), the timing of DR expression can be regulated as needed.^27,28^ These core parameters each establish a fundamental facet of DR transformation that should be pondered as the development of this technology progresses (Table 1).

**Table 1.** Applications for Optimization Parameters When Applying DRs to a New System. As DRs are applied to more plant species, optimizations specific to those species will be necessary. Following the optimization parameters we defined, which are summarized in this table, will allow for those challenges to be met.

| Optimization Parameter | Use/Application |
| --- | --- |
| DR Vector Construction | If single DR vector is less functional in species of interest, test alternate vector design schemes. |
| WUS Design | If the species specific WUS response is less effective and needs improvement, test additional variants. |
| Agrobacterium Strain | If Agrobacterium infection is not efficient, test alternate strains to find the best delivery platform for target species. |
| Shoot Progression | If induced growths do not progress beyond cell division to differentiation, test DR removal platforms. |

It is important to note that the parameters tested in these studies are not exhaustive and there will be plenty of further components for testing. Even within some of the parameters we examined there are further grounds for experimentation. The most obvious example, comes in the form of synthetically engineering the DR components involved in growth and shoot production. While the dimerization domain deletions show marked improvement in growth production, it is just one of the functional domains within *WUS* that could be manipulated.^12,19^ Certain domains are obligate for proper *WUS* function, while others are not, such as the HAM binding domain and the EAR-like domain.^29^ These less conserved domains could serve as targets for further engineering. Additionally by controlling DRs with synthetic circuits methods can be designed to not only to improve transformation efficiency but also modulate plant development related to interesting traits. Such synthetic DR engineering is still in its early days, but results such as these highlight ways in which these components can be optimized for biotechnology applications.

Expanding beyond *N. benthamiana*, the trends we have discovered to improve DR application in will likely have variable effects in different species. This can clearly be seen with the difference in growth induction potential by delivering the DRs in trans. Whereas monocot applications have seen positive effects from such delivery schema,^11^ our *trans* delivery of DRs to *N. benthamiana* seedlings was significantly less effective than the *cis* orientation (Figure 2b-c). Despite clear species distinction, certain components are universal across similar approaches. One example is the pivotal role of Agrobacterium when it is used as a delivery agent. The finding that each Agrobacterium strain presents a different growth production potential will prove to be vital for all the systems for which Agrobacterium is the primary reagent delivery platform. Knowing this principle, researchers will now have the foresight to test to determine the best Agrobacterium strain for the target tissue in their species of interest.

Each of these optimized parameters represents a small step towards developing transformation platforms that are highly scalable. It is when all these smaller optimizations are combined does the larger progress manifest towards the goal of increased efficiency. By establishing the rules through which a more functional DR based transformation platform functions, new and more efficient methods can be built across species. The advent of these more widespread transformation pipelines will thus serve as the launching point for high-throughput gene manipulation promised with technologies like CRISPR/Cas9.

## Author contributions

RAN performed the synthetic regular experiments, tested the different Agrobacterium strains and wrote the manuscript. JPC performed the vector orientation experiments, tested the CRE removal system and wrote the manuscript. MHZ performed sequence alignments to define the dimerization domain and cloned the dimerization mutant. NBB completed a portion of the growth induction co-culture experiments. A-S established the RUBY intensity code. AS performed RUBY delivery co-culture experiments. MFM cloned the developmental regulators from the alternate species. BKB funded the research. DFV funded the research and wrote the manuscript.

## Supplementary Materials

**Supplemental Table 1.**
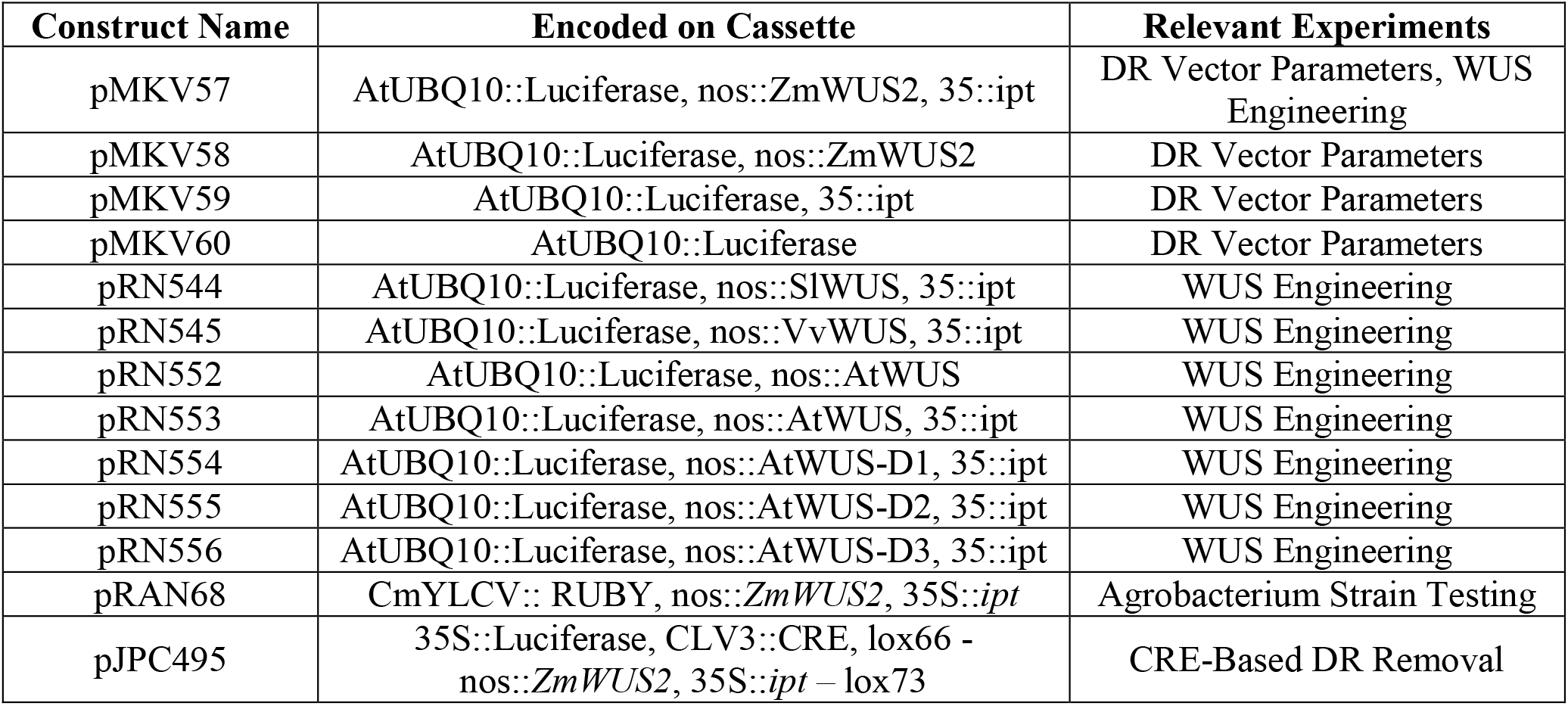
Construct List.

**Supplemental Table 2:** Growth Induction for Different Agrobacterium Strains. Strains from different labs were tested for their capacity for growth induction (growths/seedling).

| Parent Strain |  |
| --- | --- |
| GV3101 | MFM: 8.7 Voytas: 2.4 Carter: 2.1 Blackman: 1.7 |
| C58C1 | Voytas: 5.8 |
| AGL1 | Voytas: 5.6 Blackman: 2.4 |
| EHA105 | Blackman: 3.3 |

**Supplemental Figure 1.**
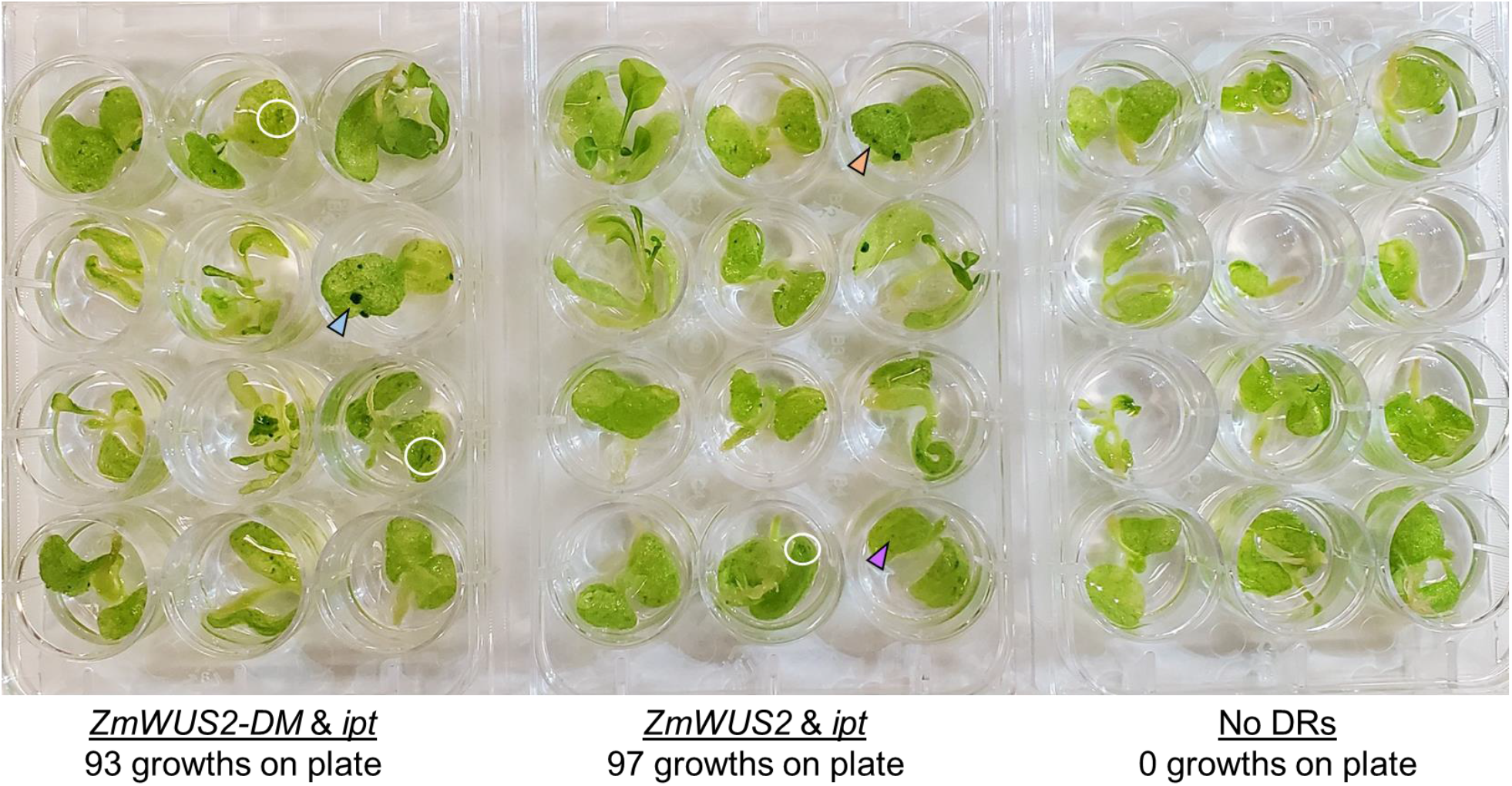
Counting and Assessing Growth Formation. Induced growths that form on *N. benthamiana* cotyledons are counted 20 days after removal from co-culture. Growths to be counted can vary in size at the time of counting. Large growths (blue arrow) are the most obvious, while more moderately sized growths (orange arrow) will make up a sizeable fraction of total growths. The hardest to define are the smaller growths that are earlier in their formation (purple arrow), which tend to look like darker green spots on the cotyledon. Growths can also form in clustered groupings (white circles), from which best growth estimates are made but counting the darker puncta within the grouping.

**Supplemental Figure 2.**
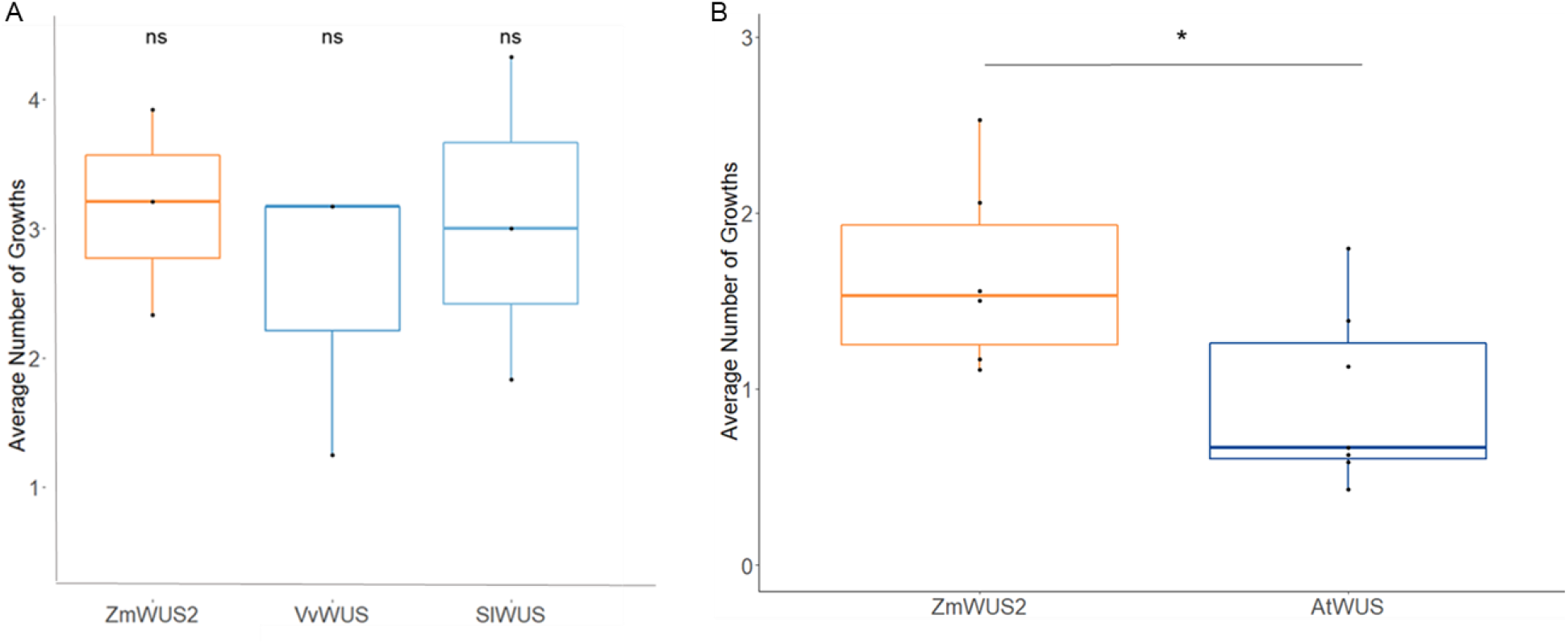
Comparing the Efficacies for *WUS* Variants from Different Species. After delivery of *WUS* variants from three different species (*ZmWUS2*, *VvWUS*, *SlWUS*) via Fast-TrACC, the number of forming growths was counted for each (a). There was no significant difference between the *ZmWUS2*, *VvWUS* and *SlWUS* variants, indicating the potential to use any of these variants for growth formation. Comparing the *ZmWUS2* variant to the *WUS* another dicot, *Arabidopsis thaliana*, it was seen that the *AtWUS* has a significantly lower pattern of growth formation (b).

**Supplemental Figure 3.**
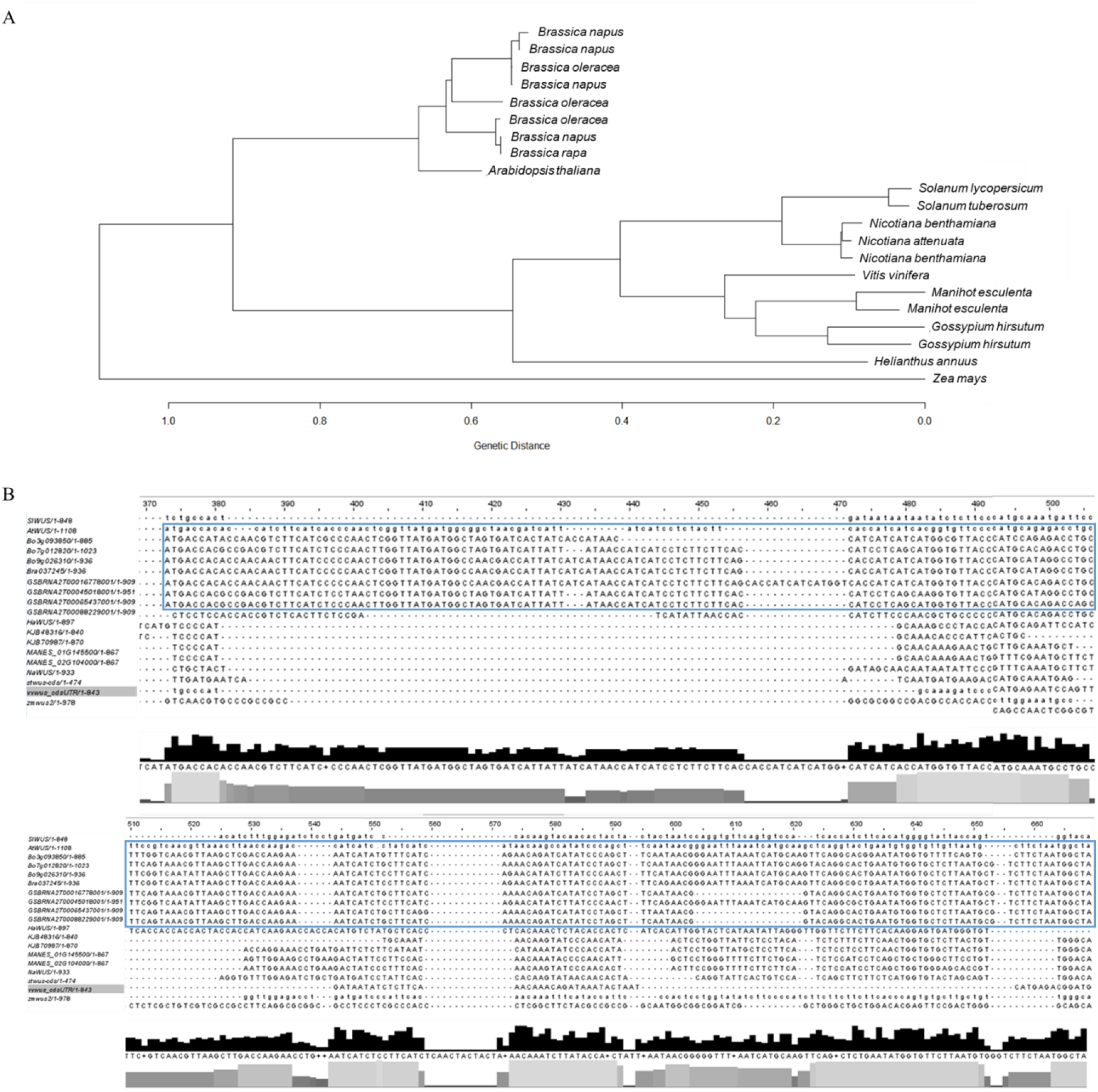
Brassica *WUS* Variants Cluster Separately from the Other Dicot *WUS* Sequences. When performing phylogenetic clustering of the *WUS* coding sequences across various different dicot species (a), interestingly all of the *WUS* variants from Brassica species clustered separately from the rest of the dicot *WUS* sequences (*ZmWUS2* serves as an out-group). This is presumably due to a conserved sequence stretch after the *WOX* homeobox domain that is highly conserved in the Brassica *WUS* sequences (b, blue box) but not in the other dicots.

**Supplemental Figure 4.**
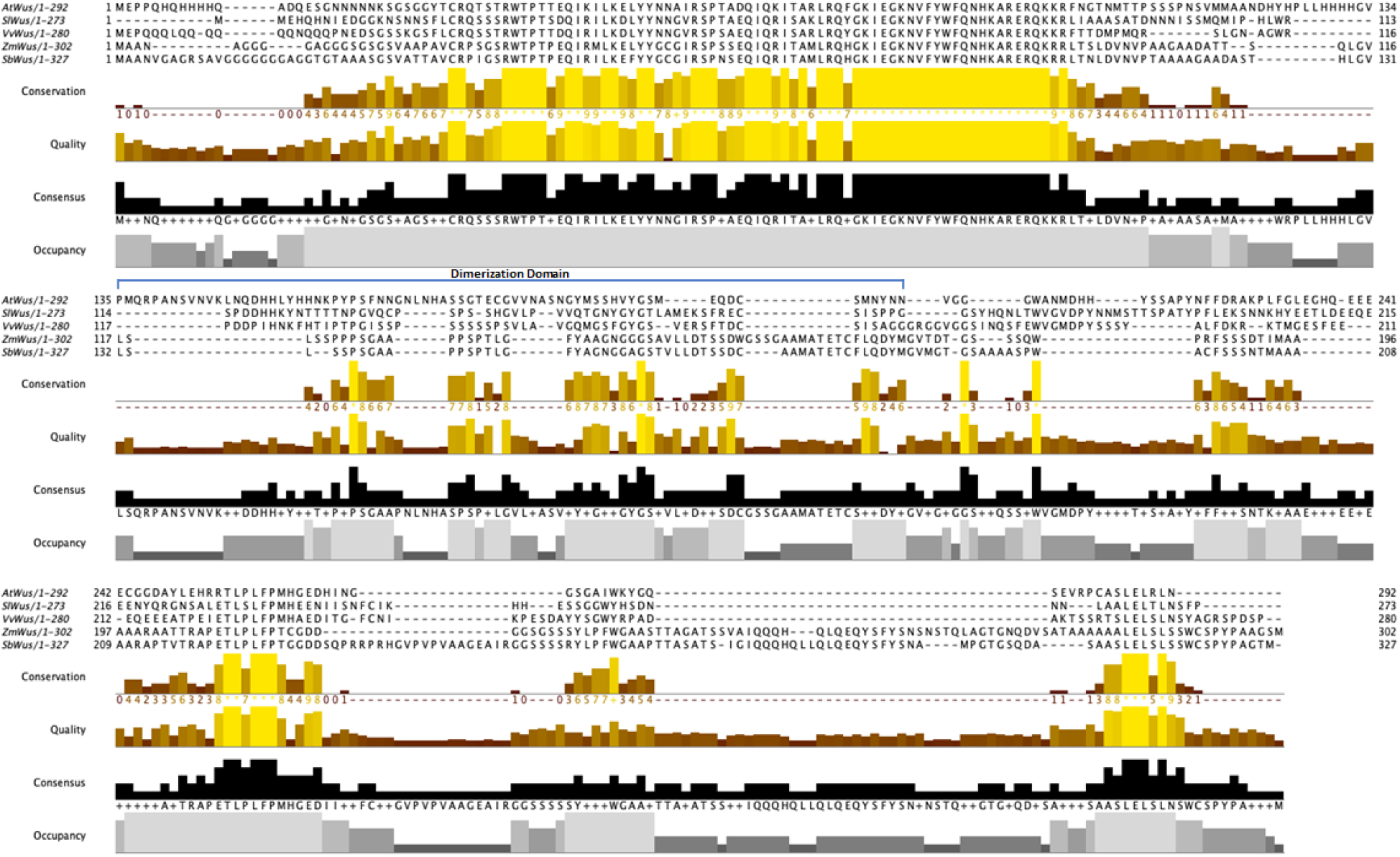
*WUS* Multiple Sequence Alignment to Define the Homodimerization Domain. WUS peptide sequences for *Arabidopsis thaliana*, *Solanum lycopersicum*, *Vitis vinifera*, *Zea mays*, and *Sorghum bicolor* were aligned using Clustal Omega. The alignment was then imported to Jalview to generate the figure shown above. Jalview scores the alignment on multiple parameters including Conservation, Quality, Occupancy, and Consensus alignment. These metrics all indicate conserved regions of *WUS* across the species. The dimerization domain is highlighted in blue and is based on the *Arabidopsis thaliana* annotation of this domain.^16^ The putative *ZmWus* truncation was determined based on the multiple sequence alignment and previously described *Arabidopsis* dimerization domain.

**Supplemental Figure 5.**
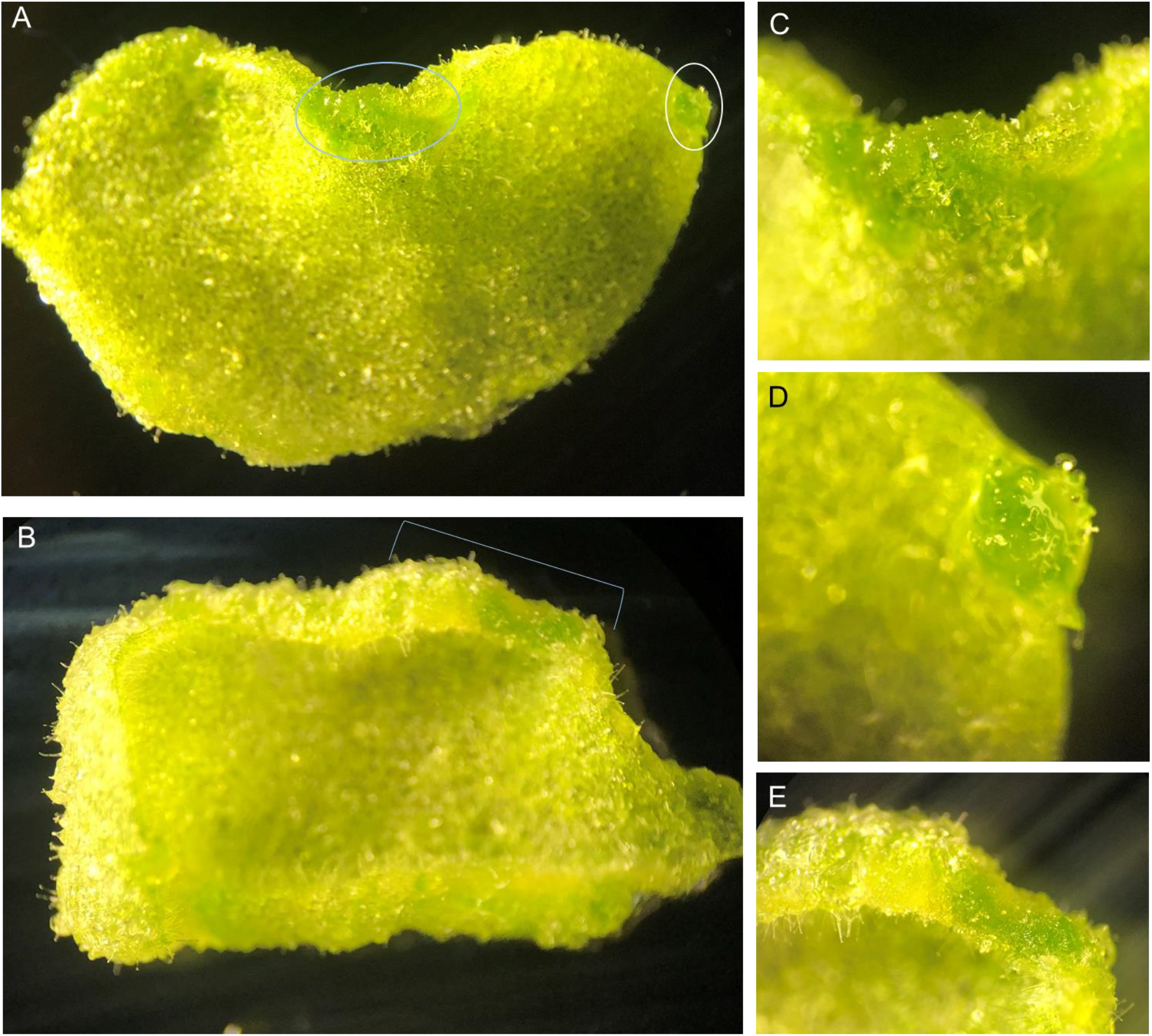
Stretches of Amorphous Growth Induced with *ZmWUS2-DM*. In addition to the individual condensed growths observed with both *ZmWUS2* variants (a: white circle, c), the *ZmWUS2*-DM variant manifests longer stretches of dedifferentiation (a-b; blue circles, d-e).

**Supplemental Figure 6.**
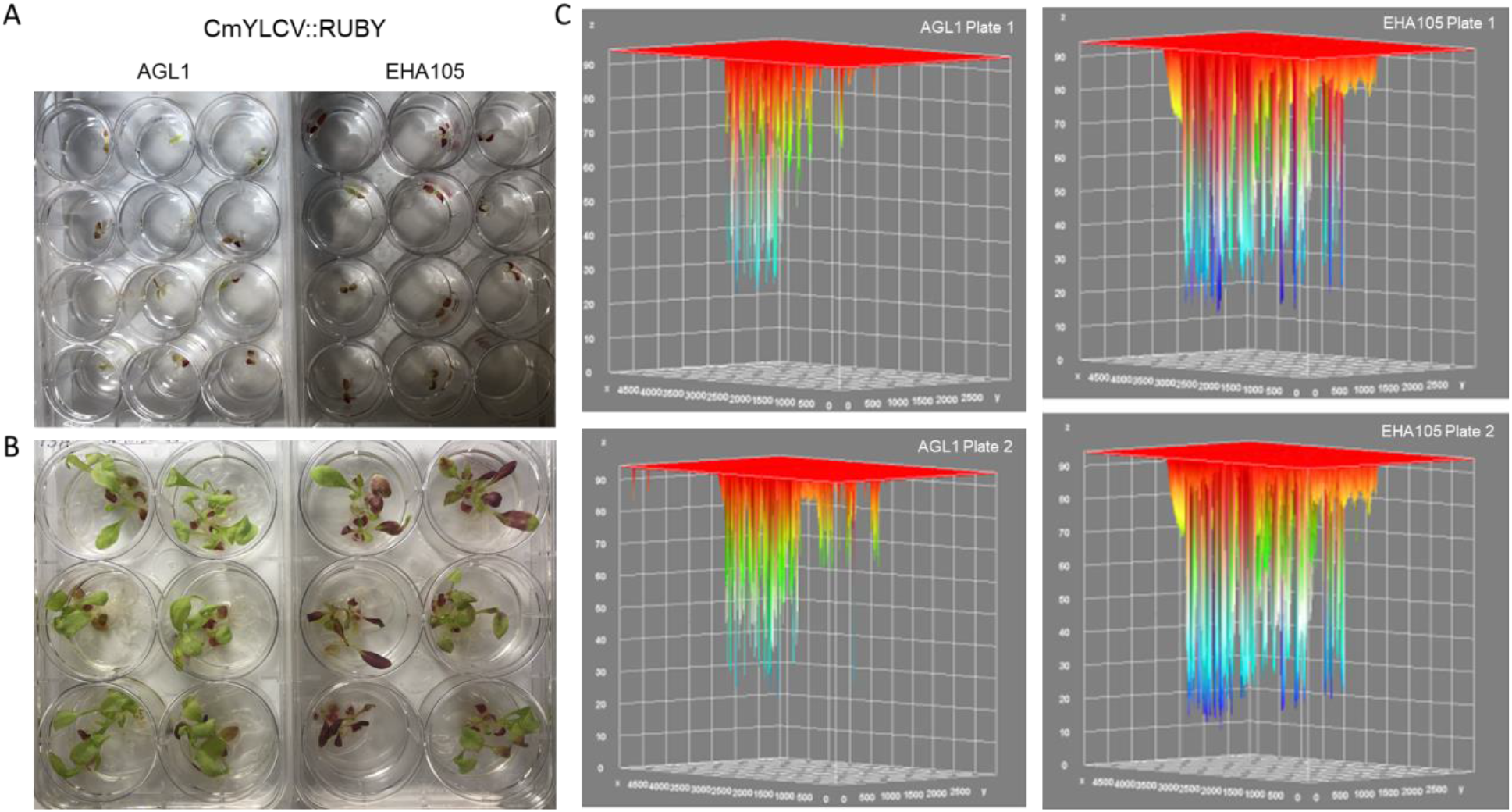
Whole Plate RUBY Intensity Differences between Agrobacterium Strains. Comparing RUBY cassette expression after Fast-TrACC delivery, initial red pigmentation seems more favored in the EHA105 seedlings (a). This difference becomes more obvious after the emergence of extant leaf primordia that were also likely transformed during the co-culture (b). Specifically at the earlier stages of testing, overall red pigmentation intensity was also stronger in the EHA105 treated seedlings (c6).

**Supplemental Figure 7.**
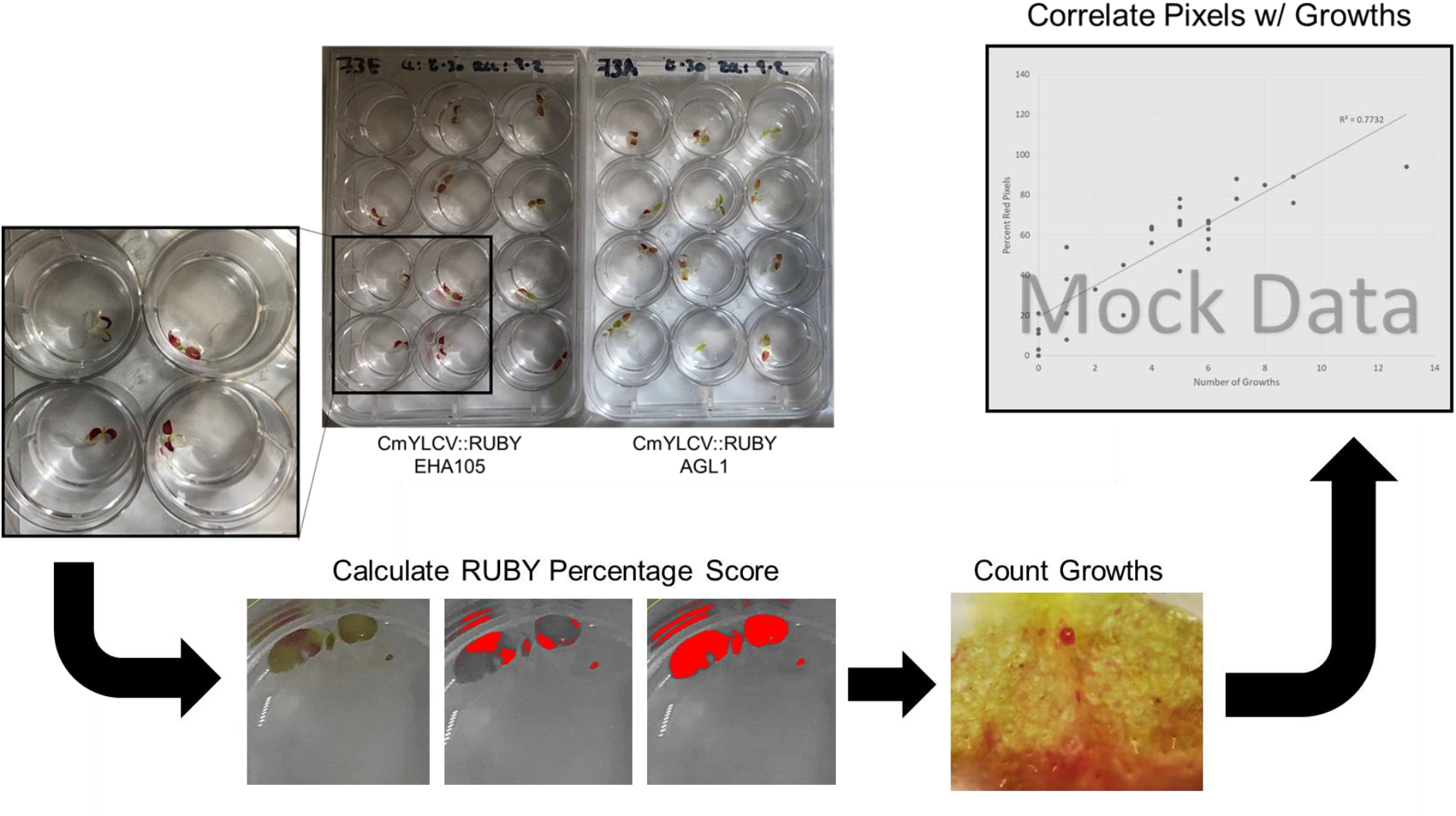
RUBY Score Calculation Pipeline. In order to determine the extent of RUBY delivery to individual seedlings, the red pigmentation was isolated in a single channel and taken as a fraction of the total seedling area. This RUBY score was then taken in comparison with the number of growths observed on that same seedling. With these two variables, a delivery and growth induction trend was made using “loess”-based statistical analyses.

**Supplemental Figure 8.**
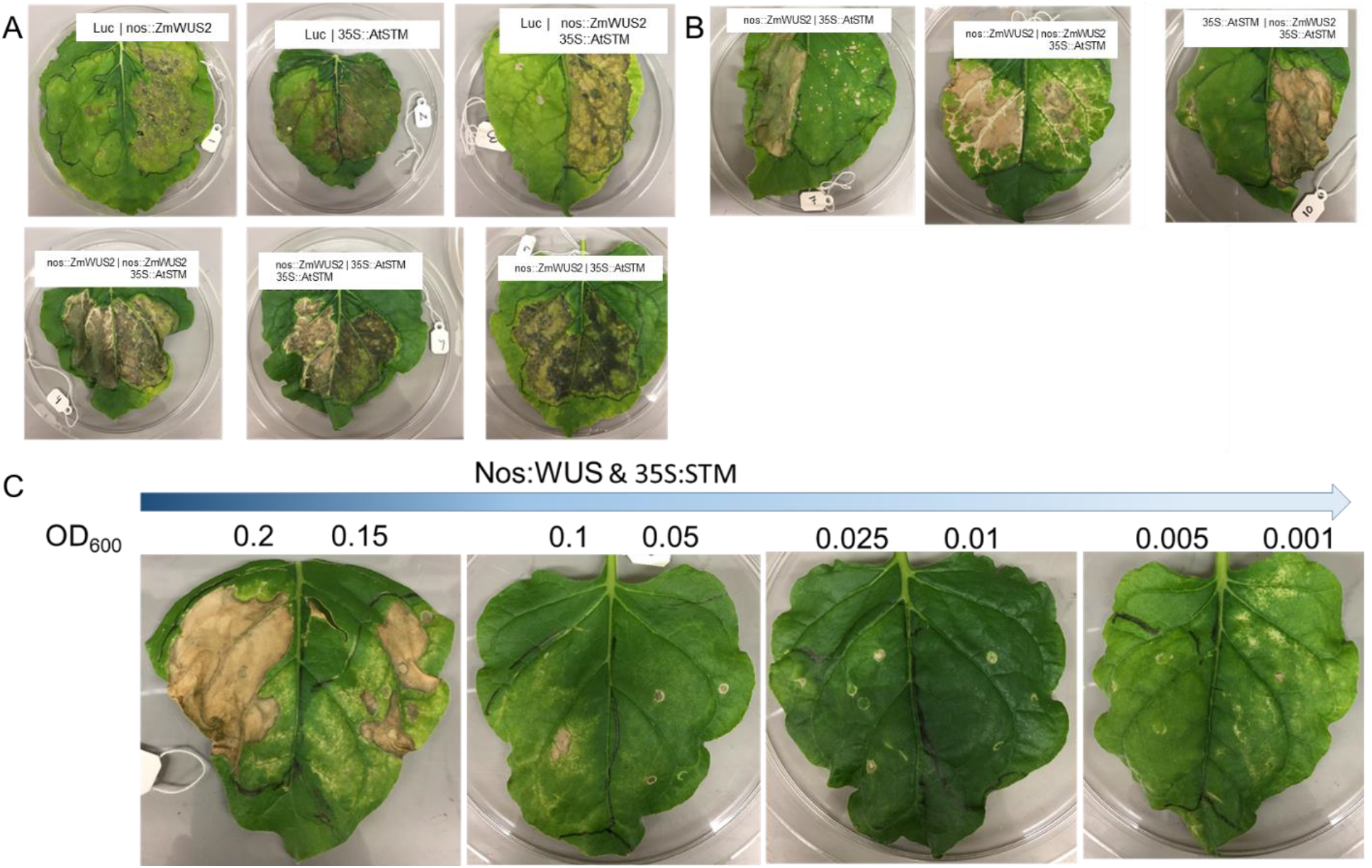
Overexpression of DRs Causes Tissue Death. Looking to test the toxicity of DR overexpression, leaf infiltation experiments were performed in *Nicotiana benthamiana* leaves. The DR cassettes tested were encoded either on geminiviral replicons (a) or standard T-DNA (b) constructs. With these infiltrations we observed a strong tissue death with *ZmWUS2* on either construct or *AtSTM* when delivered on a replicon. To mitigate this phenotype, we performed a serial agrobacterium dilution (c) and from infiltrations with lower amounts of the DRs decreased tissue death was observed.

## Notes

### Competing Interest Statement

M.F.M., R.A.N. and D.F.V. are named inventors on a patent application pertaining to the technology that was filed by the University of Minnesota. D.F.V. serves as Chief Science Officer for Calyxt, an agricultural biotechnology company that uses gene editing to create new crop varieties. All other authors have no competing interests.

